# A *Lotus japonicus* E3 ligase interacts with the Nod factor receptor 5 and positively regulates nodulation

**DOI:** 10.1101/269506

**Authors:** Daniela Tsikou, Estrella E. Ramirez, Ioanna S. Psarrakou, Jaslyn E. Wong, Dorthe B. Jensen, Erika Isono, Simona Radutoiu, Kalliope K. Papadopoulou

**Affiliations:** Department of Biochemistry and Biotechnology, University of Thessaly, Biopolis, Larisa 41500, Greece; Department of Molecular Biology and Genetics, Aarhus University, Gustav Wieds Vej, Aarhus 8000 C, Denmark; Department of Plant Systems Biology, Technical University of Munich, Emil-Ramann-Strabe 4, Freising, Germany

**Keywords:** E3 ligase, nodulation, *Lotus japonicus*, PUB13, ubiquitination, symbiosis

## Abstract

Post-translational modification of receptor proteins is involved in activation and de-activation of signaling systems in plants. Both ubiquitination and deubiquitination have been implicated in plant interactions with pathogens and symbionts. Here we present *LjPUB13*, a PUB-ARMADILLO repeat E3 ligase that specifically ubiquitinates the kinase domain of the Nod Factor receptor NFR5 and has a direct role in nodule organogenesis events in *Lotus japonicus*. Phenotypic analyses of three LORE1 retroelement insertion plant lines revealed that *pub13* plants display delayed and reduced nodulation capacity and retarded growth. *LjPUB13* expression is spatially regulated during symbiosis with *Mesorhizobium loti*, with increased levels in young developing nodules. Thus, *Lj*PUB13 is an E3 ligase with a positive regulatory role during the initial stages of nodulation in *L. japonicus*.

## INTRODUCTION

The legume-rhizobia symbiosis leads to the formation of novel organs on the plant root, termed nodules. Rhizobia within nodule cells differentiate into bacteroids that fix atmospheric dinitrogen in exchange for plant carbohydrates. The symbiotic signalling process is initiated when rhizobia secrete nodulation (Nod) factors upon sensing flavonoids produced by compatible legumes. *Lotus japonicus* Nod factor receptors NFR1 and NFR5 and the corresponding proteins LYK3 and NFP in *Medicago truncatula* (Limpens et al., 2003; Madsen et al., 2003; Radutoiu et al., 2003; Arrighi et al., 2006; Mulder et al., 2006; Smit et al., 2007) are crucial for perception of rhizobial Nod factors. Rhizobia enter into roots through infection threads (ITs) that, in most cases, initiate in epidermal root hair cells and progress to inner root tissues (reviewed in Oldroyd et al., 2011). Formation of functional nodules requires two separate but tightly coordinated developmental processes: bacterial infection and nodule organogenesis. Protein ubiquitination has been identified as being critical for these two signalling pathways.

Ubiquitination of proteins, by which proteins are tagged by ubiquitin and subsequently destined to be degraded by the proteasome, is a regulatory process essential for eukaryotic growth, development and response to interacting microbes (Vierstra, 2009). In some cases, ubiquitination is important for regulating the activity or trafficking of the target protein (Komander, 2009). Post-translational modification by ubiquitination is accomplished by a three-step process that involves ATP-dependent activation of ubiquitin by an E1 enzyme, followed by conjugation by an E2 enzyme and specific ubiquitin ligation to substrate proteins by direct interaction by an E3 ligase (Hershko and Ciechanover, 1998; Pickart and Eddins, 2004).

E3 ligases are divided into families based on their mechanism of action and on the presence of specific E2 interacting domains, such as HECT, RING and U-box (reviewed in Smalle and Vierstra, 2004; Stone and Callis, 2007). The largest class of plant U-box (PUB) proteins is the ARMADILLO (ARM) domain-containing PUB proteins that contain tandemly-repeated ARM motifs located at the C-terminal (Mudgil et al., 2004; Samuel et al., 2006). ARM repeat proteins are known to be involved in a number of different cellular processes including signal transduction, cytoskeletal regulation, nuclear import, transcriptional regulation, and ubiquitination. PUB-ARM proteins have been implicated in plant receptor-like kinase (RLK) signalling, with the ARM repeat domain mediating the binding of PUBs to the kinase domain (Gu et al., 1998; Samuel et al., 2006).

E3 ligases have been shown to play roles in the establishment of legume-rhizobium symbiosis (reviewed in Hervé et al., 2011). *M. truncatula LIN* (Kiss et al., 2009) and the orthologous gene from *L. japonicus, CERBERUS* (Yano et al., 2009), encode E3 ligases containing U-box, ARM and WD-40 repeats, and have been reported to control rhizobial infection inside root hairs. A second PUB-ARM E3 ubiquitin ligase in *M. truncatula, Mt*PUB1, was identified as a negative regulator of nodulation by direct interaction with the receptor-like kinase LYK3 (Mbengue et al., 2010). *M. truncatula* PUB1 is required for both rhizobial and arbuscular mycorrhiza (AM) endosymbiosis as it also directly interacts with the receptor kinase DMI2, a key component of the common symbiosis signalling pathway (Vernié et al., 2016). A member of the SEVEN IN ABSENTIA (SINA) family of E3 ligases, SINA4, was shown to interact with SYMRK receptor-like kinase in *L. japonicus* and be a negative regulator of rhizobial infection (Den Herder et al., 2012). *Lj*nsRING, a RING-H2 E3 ubiquitin ligase from *L. japonicus*, was also reported to be required for both rhizobial infection and nodule function (Shimomura et al., 2006). Nevertheless, the mechanistic mode of action of the E3 ligases in the symbiotic interactions has not been elucidated.

Here, we report the involvement of a PUB-ARM protein, *Lj*PUB13, in the symbiotic interaction of *L. japonicus* with rhizobia. *Lj*PUB13 is phylogenetically a close relative of *Arabidopsis* PUB13, which has been implicated in plant responses to bacterial flagellin (flg22) (Lu et al., 2011) and, very recently, fungal long-chain chitooligosaccharides (chitooctaose) (Liao et al., 2017). In *L. japonicus, Lj*PUB13 is involved in the establishment of a successful symbiosis, through the interaction of its ARM domain with the Nod factor receptor NFR5 and direct ubiquitination of the NFR5 kinase domain.

## MATERIALS AND METHODS

### Biological material, growth conditions and inoculation

*L. japonicus* ecotype Gifu B-129 was used as a wild type control. Wild type, *pub13* and *har1-3* (Krusell at al., 2012) seeds were surface-sterilized as previously described (Handberg & Stougaard, 1992) and were grown on wet filter paper for 3 to 4 days. Plants were then grown in petri dishes with solid quarter-strength B&D medium (Broughton & Dilworth, 1971) on filter paper. The plants were grown in a vertical position in growth boxes, keeping the roots in the dark. Growth chamber conditions were 16-h day and 8-h night cycles at 21°C.

For nodulation kinetics and IT counting, each petri dish was inoculated with 500 μl of a 0.02 OD_600_ culture of *Mesorhizobium loti* cv. R7A DsRed (Kawaharada et al., 2015). Nodule numbers were scored on at least 30 plants and ITs were counted on at least 30 cm of root.

For gene expression analyses the plants were grown in pots with 2:1 mix of sand and vermiculite. Half of the plants were inoculated with a 0.1 OD_600_ culture of *M. loti* cv. R7A. The plants were watered periodically with Hoagland nutrient solution.

### Protein purification

For GST-*Lj*PUB13, GST-*Lj*BAK1_cyt_, and GST-*Lj*FLS2_cyt_, the corresponding ORFs were PCR amplified and the resulting fragments were cloned between the EcoRI and XhoI sites of pGEX-6P-1 (GE Healthcare). For HIS-*Lj*PUB13, the ORF was PCR amplified and the resulting fragment was cloned between the EcoRI and XhoI sites of pET21a (Novagen). Primers are listed in Table S1. GST-and HIS-tagged proteins were expressed in *E. coli* Rosetta (DE3) (Merck Chemicals). GST-tagged proteins were then purified with Glutathione-Sepharose 4B beads (GE Healthcare) and eluted with 0.2 M reduced glutathione, whilst HIS-tagged proteins were purified with Ni Sepharose High Performance beads (GE Healthcare) and eluted with 250 mM imidazole.

Plasmids with the *NFR1*_cyt_ and *NFR5*_cyt_ fragments cloned in pProEX-1 vector were kindly provided by Mickael Blaise. HIS-NFR1_cyt_ and HIS-NFR5_cyt_ proteins were expressed in Rosetta 2 *E. coli* (DE3) competent cells (Novagen). IMAC purification was performed using Ni-NTA columns (Qiagen). The proteins were eluted with an elution buffer containing 50 mM Tris-HCl pH 8, 500 mM NaCl, 500 mM imidazole pH 8, 1 mM benzamidine, 5 mM β-mercaptoethanol and 10% glycerol. The eluted proteins were then injected onto a Superdex 200 increase 10/300 GL (GE Healthcare) column connected to an ÄKTA PURIFIER system (GE Healthcare) and eluted with a buffer containing 50 mM Tris-HCl pH 8, 500 mM NaCl, 5 mM β-mercaptoethanol and 10% glycerol.

### *In vitro* ubiquitination assay

The *in vitro* ubiquitination assays were performed with an Ubiquitinylation kit (BML-UW9920, Enzo Life Sciences), using either UbcH6 or UbcH5b E2 enzymes and following the manufacturer’s protocol. For *Lj*FLS2 ubiquitination tests UbcH5c E2 enzyme was also used.

All the reactions were incubated at 30°C for 3 hrs, and then stopped by adding SDS sample buffer and boiled at 98°C for 5 min. The samples were then separated by SDS–PAGE and analyzed by Western blotting, using the a-GST antibody (GE Healthcare) and the a-HIS antibody (Roche) to detect the tagged proteins or the anti-ubiquitin antibody (Santa Cruz Biotechnology P4D1) to detect the ubiquitinated fraction.

### *In vitro* binding assay

For the *in vitro* binding assays, 2 μg of a GST tagged protein was incubated with 2 μg of a HIS tagged protein, in different protein combinations, in 150 μl of cold buffer A (50 mM Tris-HCl pH 7.5, 100 mM NaCl, 10% glycerol) with 0.1 % Triton X-100, for 1.5 hrs at 4^°^C. Glutathione-Sepharose 4B beads (GE Healthcare) were added to the mixtures and incubated with the proteins for 2 more hrs at 4^°^C. The beads were then washed 3 times with cold buffer A. The bead-bound proteins were analysed by immunoblotting, according to standard protocols, using anti-GST (GE Healthcare) and anti-HIS (Roche) antibodies.

### Expression analysis by qRT-PCR

To test the temporal and spatial expression of *LjPUB13* in non-inoculated and *M. loti* inoculated *L. japonicus* plants and the expression of defence genes after treatment with flg22, analysis by qRT-PCR was performed as previously described (Kawaharada et al., 2015; Tanou et al., 2015). Gene primers are listed in Supplementary Table S2.

### *LjPUB13* Promoter Activity in *L. japonicus*

For the promoter-GUS-terminator construction, a 1475 bp promoter with a 5′untranslated region (UTR) and a 310 bp terminator region was amplified from *L. japonicus* genomic DNA (the primers are listed in Table S3) and cloned into GoldenGate vectors (Weber et al., 2011). The PCR fragments were firstly cloned into GoldenGate Level 0 vectors before being assembled as a construct (promoter:GUS:terminator) in a modified pIV10 vector (Stougaard et al., 1987). The construct was transferred into *A. rhizogenes* AGL1.

Hairy root induction using *A. rhizogenes* was performed as described previously (Hansen et al., 1989). Chimeric plants were transferred into magenta growth boxes containing a sterilized 4:1 mix of clay granules and vermiculite as well as ¼ strength B&D medium supplemented with 1 mM KNO_3_. For inoculation, liquid cultures of *M. loti* cv. R7A expressing DsRed were grown to an optical density of 0.02 and applied directly to the root systems (0.7 ml per plant). Plants were grown at 21°C (16 h light, 8 h dark) and harvested at indicated times post inoculation. GUS staining was performed as described previously (Vitha et al., 1995). Whole roots were visualized on a Leica M165 FC stereomicroscope.

### Infection thread formation

For inspection of infection-thread formation, roots were harvested 10 and 14 days after inoculation with *M. loti* strain cv. R7A DsRed. Sections (1 cm) of at least 30 plants were examined under a Zeiss Axioplan 2 fluorescent microscope.

### ROS accumulation

Seven-day-old roots were cut into 0.5 cm pieces and incubated overnight (with shaking) in 200 µl water in 96-well plates (Grenier Bio-one). Before the measurements, the water was exchanged with 200 µl of buffer (20 mM luminol, Sigma; 5 µg/ml horseradish peroxidase, Sigma), supplemented with either H_2_O or 0.5 µM flg22 peptide (FLS22-P-1, Alpha Diagnostic). Luminescence was recorded with a Varioskan™Flash Multimode Reader (Thermo).

### Protein-protein interaction studies by Bimolecular Fluorescence complementation (BiFC)

The ARM domain of *LjPUB13* (Fig. S2) and the full length *LjBAK1* were cloned, using Gateway technology (Invitrogen), into the pGREEN029:35S:GW:nYFP/cYFP vectors creating N- and C-terminal fusions to YFP. The primers are listed in Table S3. The NFR1 and NFR5 fused to nYFP/cYFP constructs used were the same as those described in Madsen et al., 2011.

*Agrobacterium tumefaciens* AGL1 cells transformed with the protein expression plasmids were grown in 5 ml LB medium supplemented with appropriate antibiotics at 28°C. Bacteria were pelleted by centrifugation at 4000 *g* for 20 min at room temperature and re-suspended in agroinfiltration medium (10 mM MgCl_2_, 10 mM MES and 450 μM acetosyringone), incubated for 2-3 hrs in the dark and finally re-suspended to an OD_600_ of 0.2. A 1:1 mixture of cultures was prepared for each construct combination together with the P19 construct (to an OD_600_ of 0.02). A 1-ml syringe was used for infiltration of the bacterial mixture to the abaxial side of the *Nicotiana benthamiana* leaves. Fluorescence was detected using a Zeiss LSM510 confocal microscope.

### Statistical analyses

Differences in the tested biological parameters between mutant and wild type plants were analyzed by Student’s t-test. A significant level of 5% was applied.

## ACCESSION NUMBERS

The *LjPUB13* sequence is available in GenBank under the accession number KY131979 and in *Lotus* base v.3.0 (https://lotus.au.dk/) as Lj3g3v3189730.1. Other sequence data from this article can be found in the GenBank under the following accession numbers: *LjBAK1* (KY131980), *LjFLS2* (JN099749), *NFR1* (AJ575249), *NFR5* (AJ575255)

## RESULTS

### *LjPUB13* encodes an active E3 ligase

A *Lotus japonicus* sequence encoding for a novel ARMADILLO repeat-containing protein was initially identified as the TC63883 in the DFCI Gene Index Database (http://compbio.dfci.harvard.edu/tgi/tgipage.html), while searching for ARMADILLO repeat proteins in *L. japonicus.* Initial search at transcript data in *Lotus* Base (https://lotus.au.dk) showed that this gene is expressed in nodules. The predicted protein has a typical Plant U-box (PUB) E3 ubiquitin ligase sequence with a U-box motif followed by a C-terminal ARMADILLO (ARM) repeat domain (Fig. 1A). It encodes a 671 aminoacid long protein with an estimated molecular mass of 73kDa. Phylogenetic analysis showed that this protein is distinct from other previously characterised PUBs in legumes, but it is the closest homolog to *Arabidopsis thaliana* PUB13 (Fig. S1, S2). We therefore designated the protein as *Lj*PUB13. The encoded polypeptide shares 69% and 65% amino acid identity with *At*PUB13 (AT3G46510) and *At*PUB12 (AT2G28830), respectively. At the genomic level, this gene displays an exon-intron structure similar to *AtPUB13* (Li *et al.*, 2012), but different from *AtPUB12* (www.arabidopsis.org), with four exons spanning a 4547 bp region on chromosome 3 (Fig. 1A).

**Figure 1.**
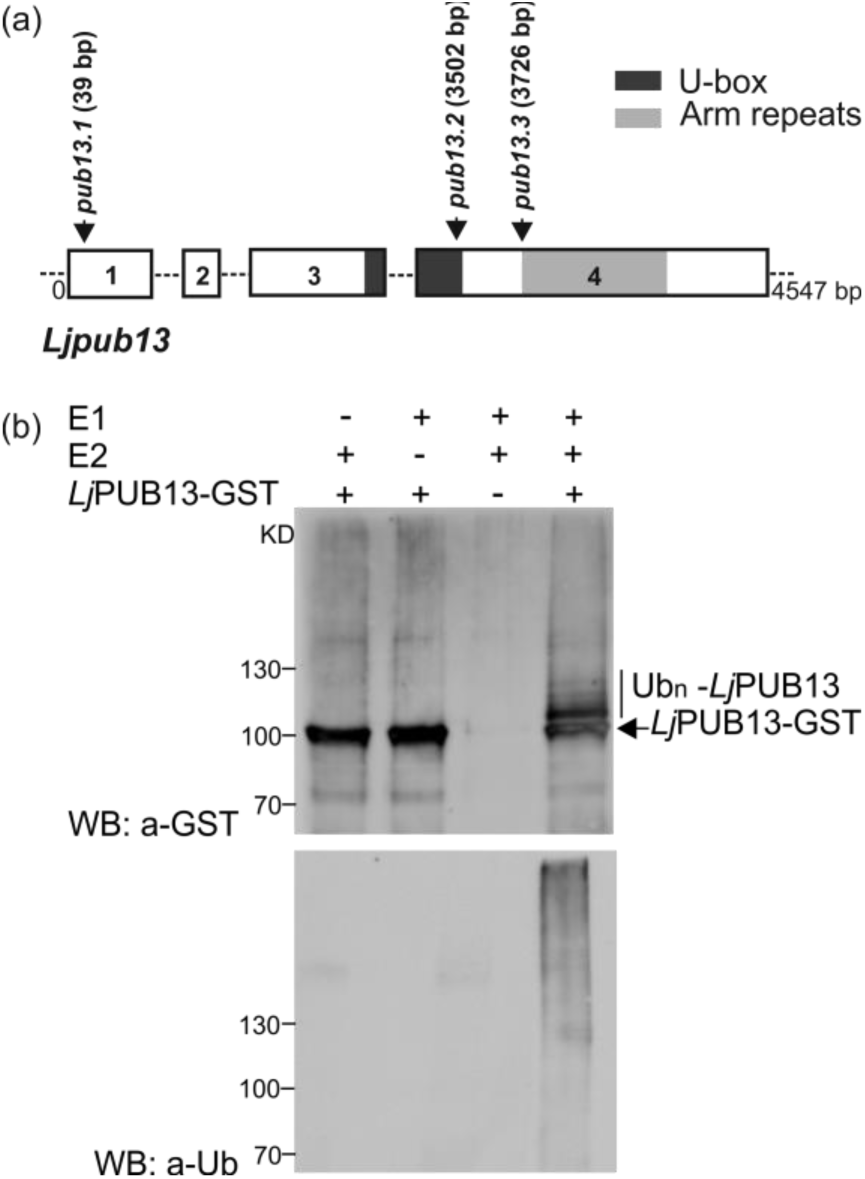
*Lj*PUB13 is an active E3 ubiquitin ligase. (**a**) *L. japonicus PUB13* gene structure. PUB13 protein contains a U-box motif and an ARM-repeat domain. The *pub13* mutants carry LORE1 insertions in the coding region of the *PUB13* gene. The positions of the LORE1 insertions in the *pub13* mutants are marked by arrows. Numbers indicate nucleotides. (**b**) *Lj*PUB13 auto-ubiquitination. PUB13 was purified as a GST fusion protein. The ubiquitination was detected by both anti-GST and anti-Ub antibodies. UbcH6 was used as the E2 enzyme in the reactions. The experiment was repeated four times with similar results.

The predicted E3 ligase enzymatic activity of *Lj*PUB13 was analysed by an *in vitro* ubiquitination assay, performed with *Lj*PUB13 purified as a glutathione S-transferase (GST) fusion protein from *Escherichia coli*. Incubation of *Lj*PUB13 with E1 ubiquitin-activating and E2 ubiquitin-conjugated enzymes, ubiquitin and ATP resulted in *Lj*PUB13 polyubiquitination (Fig. 1B). Bands with a larger molecular weight than *Lj*PUB13-GST and a protein ladder were detected by anti-GST and anti-ubiquitin antibodies, respectively, when *Lj*PUB13 was added to the reaction (Fig. 1B, lane 4). The observed E3 ligase activity was abolished in the absence of E1 or E2 enzymes (Fig. 1B, lanes 1 & 2). These results show that *L. japonicus* PUB13 is a functional E3 ligase, possessing auto-ubiquitination activity.

### The expression of *LjPUB13* is symbiotically regulated

To test if *LjPUB13* is regulated during symbiosis, we monitored its transcript levels in roots and nodules at 7, 14, 21 and 28 days post inoculation (dpi) with *Mesorhizobium loti*. No significant increase was detected in inoculated plants compared to uninoculated ones. Functional nodules, in general, had approximately 2 times less *LjPUB13* transcript compared to roots (Fig. 2A).

**Figure 2.**
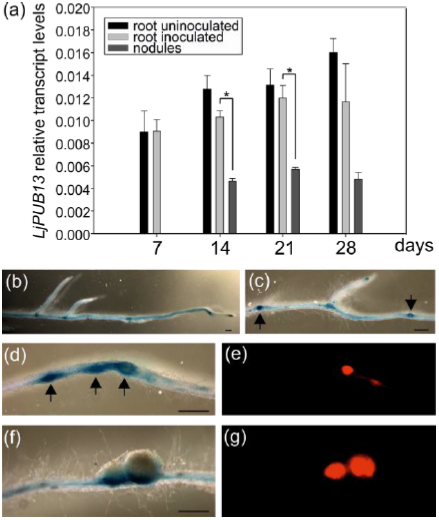
*LjPUB13* is expressed in symbiotically active root and nodules. (**a**) *LjPUB13* expression in uninoculated *L. japonicus* roots *vs* roots inoculated by *M. loti* at 7, 14, 21 and 28 days post inoculation and 14, 21 and 28-day-old nodules. Transcript levels were normalized to those of *UBQ*. Bars represent means (+SE) of three biological replications (n=8). Significant differences (*P*<0.05) are indicated by asterisk. (**b, c, d, f**) Expression of *LjPUB13* in *L. japonicus* transgenic hairy roots transformed with a *ProPUB13*::*GUS* construct, detected after GUS staining. In both uninoculated (b, c) and inoculated roots (d, f), *PUB13* promoter activity was observed in the vascular bundle. At uninoculated plants strong *LjPUB13* expression is detected in lateral root initiation sites (c, arrows). After rhizobial inoculation (d, f), *LjPUB13* promoter was active in nodule promordia and young developing nodules at 7dpi (d, arrows) but not in mature nodules at 21dpi (f). (**e and g)** show the tissues colonised by DsRed-expressing rhizobia in (d) and (f), respectively. Bars, 500 μm.

To complement the results from gene transcript analysis, we constructed a *LjPUB13*_*pro*_::GUS fusion and analysed the spatial and temporal regulation of *LjPUB13* promoter activity in *Agrobacterium rhizogenes* transformed roots expressing the transgene. In both uninoculated and inoculated roots, the *LjPUB13* promoter was active throughout the whole root, but strongest in the vascular bundle (Fig. 2B) and at the lateral root initiation sites (Fig. 2C). In *M. loti* inoculated plants, an increase in the promoter activity in the root zones, where nodule primordia formation occurs, was observed (Fig. 2D). Although strong *LjPUB13*_*Pro*_::GUS expression was detected in young developing nodules at 7 dpi (Fig. 2D), in the fully developed mature nodules (28 dpi), *LjPUB13*_*Pro*_::GUS expression was restrained to the nodule-root connection zone (Fig. 2F). Collectively, these results illustrate that *LjPUB13* transcription is present in symbiotically active root and young nodule tissues.

### *Ljpub13* mutants display growth defects and a delayed and reduced nodulation capacity

To investigate whether *Lj*PUB13 plays a role in the interaction between *L. japonicus* and the *M. loti* symbiont, we identified three mutant lines that possess LORE1 retroelement insertions (Fukai *et al.*, 2012; Urbanski *et al.*, 2012) in *LjPUB13* (Fig. 1A) and homozygous insertion lines were obtained for phenotypic analyses. The expression levels of *LjPUB13* in uninoculated plants of *pub13.1, pub13.2* and *pub13.3* mutant lines are 4-, 4.5- and 19-fold reduced, respectively, compared to wild-type plants of the same age (Fig. S3).

We observed that in the absence of the *M. loti* symbiont, *L. japonicus pub13* mutants displayed a reduced growth phenotype; the root and shoot length are significantly shorter in all *pub13* mutants when compared to wild type plants of the same age (Fig. 3). In the presence of *M. loti, pub13* mutants formed a lower number of nodules compared to the wild type Gifu, both in absolute number (Fig. 4A) and when normalized to root length (Fig. 4B). The latter indicates that the reduced nodulation phenotype does not correlate to the shorter root phenotype and is further supported by the direct comparison of *pub13.1* with the *har1* hypernodulation mutant (Krusell at al., 2012; Nishimura et al., 2002), where in *har1* very short roots (shorter than *pub13.1*) a high number of nodules is formed (Fig. S4). Notably, nodulation was not only significantly reduced but also delayed in *pub13* mutants (Fig. 4C, Fig. S5). At 14 dpi only 15% of *pub13-1* and *pub13-2* plants and 22% of *pub13-3* plants had initiated symbiosis and had nodules compared to approximately 60% of wild type plants. This frequency in nodule appearance in the different plants of each plant line was higher at later stages of nodulation; at 28 dpi 50% of *pub13-1* and almost 80% and 90% of *pub13-2* and *pub13-3* plants, respectively, formed at least 1 nodule (Fig. 4C). Sections of mature nodules showed that they were fully infected and morphologically normal (Fig. S6). In addition, the root infection process was not evidently perturbed in the *L. japonicus pub13* mutants since both wild type plants and *pub13* mutants formed a similar number of infection threads in root hairs (Fig. 4D). Together, these results show that *Lj*PUB13 is involved in plant growth and nodule organogenesis.

**Figure 3.**
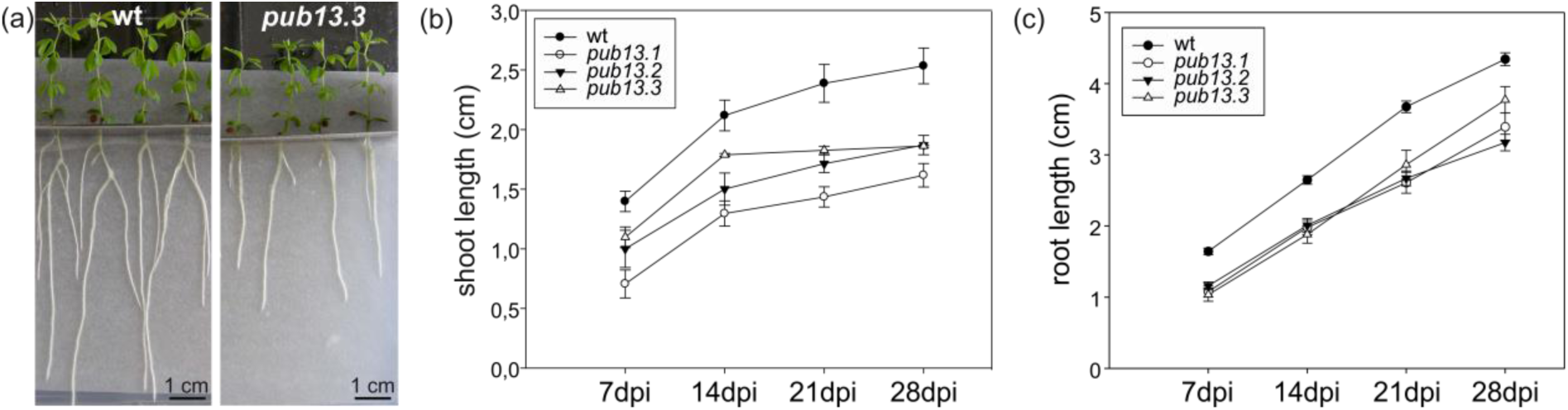
*L. japonicus pub13* mutants have a reduced growth phenotype. (**a**) *L. japonicus pub13.3* plants have shorter roots and shoots compared to wild type. (**b**) Shoot length and (**c**) root length of *pub13* mutants is significantly shorter compared to wild type plants. Graphs show means (+SE) of three biological replications (n=10). (b, c) The differences between wt and *pub13* mutants are statistically significant at all time points (P=0.001; one-way ANOVA and Tukey test), except wt-*pub13.3* shoots at 7 and 14 dpi.

**Figure 4.**
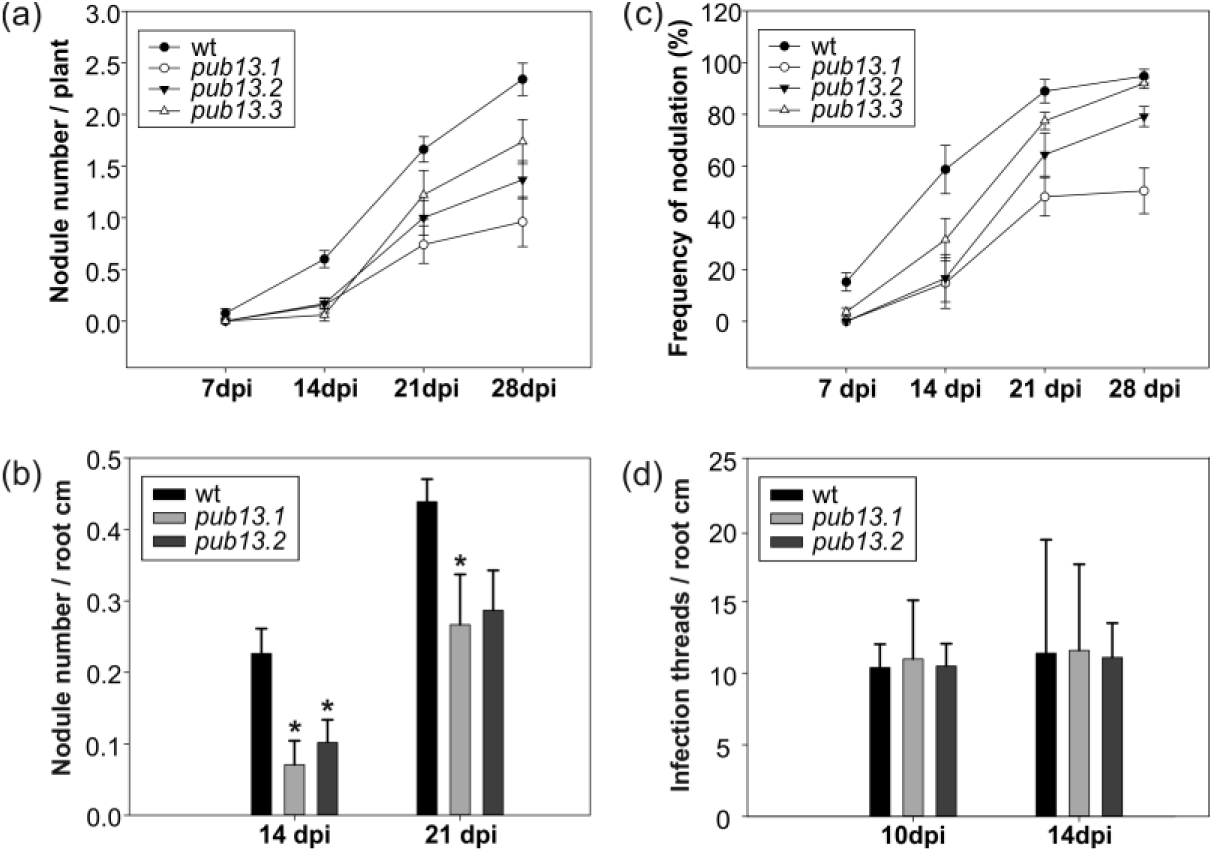
Mutants of *LjPUB13* have reduced nodulation. (**a**) Nodulation kinetics of wild type and *pub13* mutants, (**b**) number of nodules per root cm in *pub13.1* and *pub13.2* mutants compared to wild type plants at 14 and 21 dpi, (**c**) frequency of plants carrying nodules, (**d**) number of infection threads per root cm in *pub13.1* and *pub13.2* mutants compared to wild type plants at 10 and 14 dpi. Graphs show means (+SE) of at least three biological replications (n=10). (a, c) the differences between wt and *pub13* mutants are significant (P<0.05; one-way ANOVA and Tukey test) at all time points, except wt-*pub13.3* at 21 and 28 dpi.

### *L. japonicus pub13* mutants display normal root PTI responses to flg22 treatment

Based on its gene structure, amino acid identity and phylogenetic relationship, *LjPUB13* could be considered a putative ortholog of *AtPUB13* in *L. japonicus.* In *Arabidopsis*, a number of phenotypes have been reported for *Atpub13* mutants, which primarily indicates for a role in PTI (pathogen-associated molecular patterns (PAMP)/pattern-triggered immunity) response to flg22 treatment (Lu et al., 2011, Zhou et al., 2015). Although the studies in *Arabidopsis* were focused in leaves, we searched for similar functions in *L. japonicus* roots as our aim was to investigate the function of *LjPUB13* in root responses to interacting microbes or microbial signals.

Firstly, we investigated the involvement of *Lj*PUB13 in the induction of reactive oxygen species (ROS) after flg22 treatment. Our analysis of wild type and *pub13* mutant roots revealed that similar levels of ROS were produced by wild type and mutant *L. japonicus* plants after treatment with 0.5 μM flg22 (Fig. S7).

Secondly, we analysed the transcriptional activation of defence marker genes *LjMPK3, LjPEROXIDASE* and *LjPR1*, together with *LjFLS2* and *LjBAK1* after 1-hour treatment with flg22. All marker genes were induced in the treated samples (Fig S8); *LjMPK3, LjPEROXIDASE* and *LjPR1* had a 4- to 6-fold increase in transcript levels while a 2- to 3-fold increase was observed for *LjFLS2* and *LjBAK1.* Nevertheless, the expression levels of these marker genes were comparable in wild type and *pub13* mutants in both treated and non-treated samples (Fig. S8), suggesting that *L. japonicus pub13* mutants neither express immunity-related genes constitutively nor overexpress them in the presence of flg22 as seen in *Arabidopsis* mutants (Lu et al., 2011).

Collectively, these results show that in *L. japonicus* roots the *PUB13* gene is not directly involved in PTI responses induced by flg22.

### *Lotus* PUB13 interacts with *Lj*BAK1 but fails to ubiquitinate *Lj*FLS2

It is known that in *Arabidopsis*, PUB13, phosphorylated by BAK1 in the presence of flg22, polyubiquitinates the FLS2 receptor. This results in FLS2 degradation and regulation of the PTI signalling downstream of FLS2-BAK1 (Lu et al., 2011). The apparent absence of PUB13-dependent PTI responses in *L. japonicus* roots prompted us to investigate the molecular basis of this differential response in *L. japonicus*.

We identified the corresponding *L. japonicus* BAK1, and the *Lj*BAK1_cyt_-GST or *Lj*FLS2_cyt_-GST fusion proteins were produced in *E. coli*. Next, we investigated the ability of the *Lj*PUB13-HIS protein to interact with *Lj*BAK1 by performing an *in vitro* binding assay. (Similar to *Arabidopsis* (Lu et al., 2011), we observed that *Lj*PUB13 interacts with *Lj*BAK1, while the *Lj*PUB13-*Lj*FLS2 interaction is barely detectable (Fig. 5A). *Lj*PUB13 appears to not to be able to ubiquitinate *Lj*FLS2_cyt_, although we tested potential capacity in the presence of three different E2 enzymes (Fig. S9). These results indicate that PUB13 has different molecular capacities in *L. japonicus* and *Arabidopsis.*

**Figure 5.**
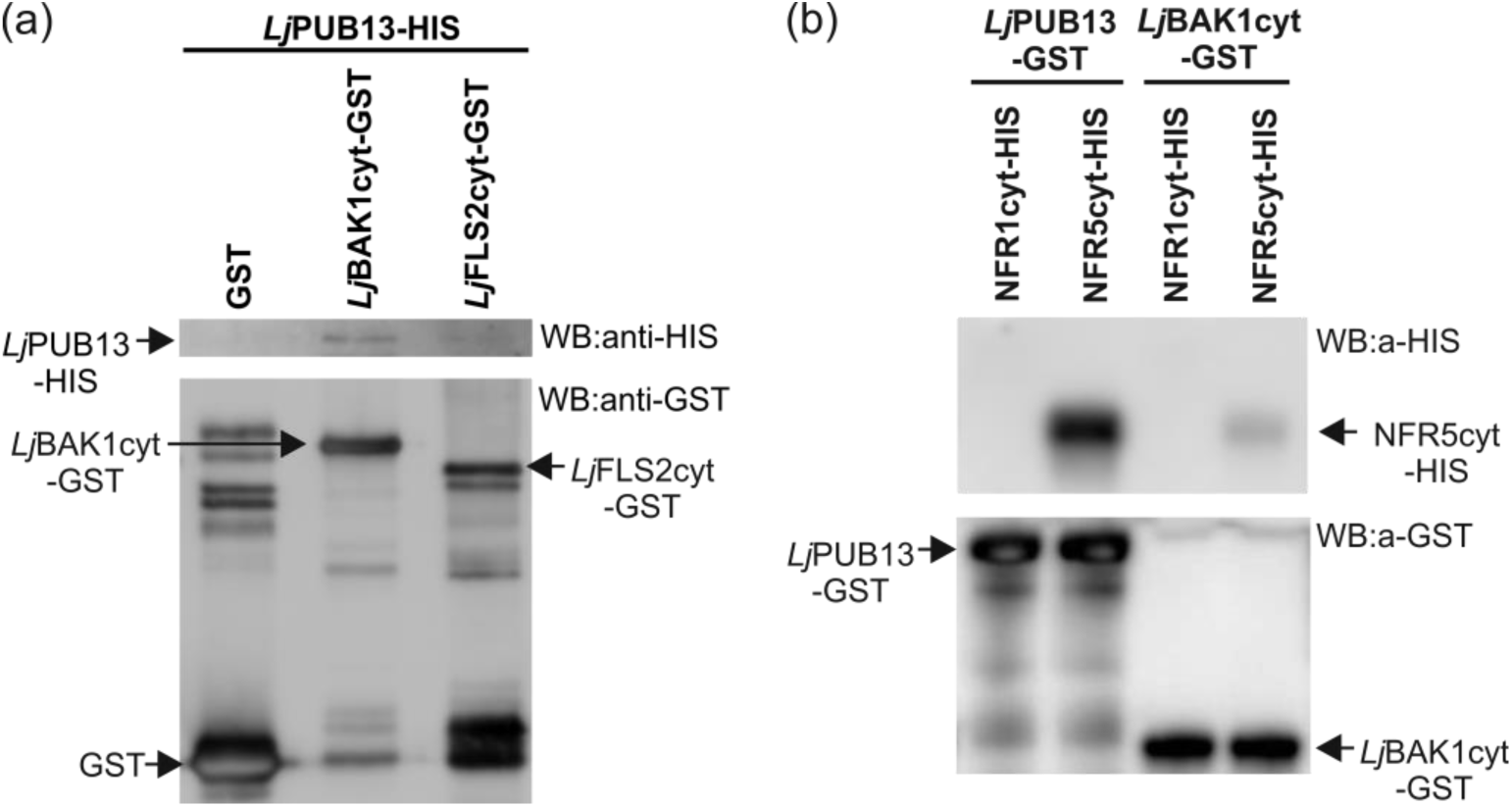
*Lj*PUB13 interacts with *Lj*BAK1, *Lj*FLS2 and NFR5 *in vitro*. **(a)** *In vitro* binding assay of the HIS-tagged *Lj*PUB13 with the GST-fused cytoplasmic regions of *Lj*BAK1 and *Lj*FLS2. *Lj*PUB13 interacts with *Lj*BAK1_cyt_ and *Lj*FLS2_cyt_. **(b)** *In vitro* binding assays of the GST-fused *Lj*PUB13 and *Lj*BAK1_cyt_ with the HIS-tagged cytoplasmic regions of the Nod Factor receptors NFR1 and NFR5. *Lj*PUB13 interacts strongly with NFR5_cyt_, while the *Lj*PUB13-NFR1_cyt_ interaction is undetectable. Sometimes a weak interaction of *Lj*PUB13-NFR1_cyt_ was observed but this result was not always reproducible. *Lj*BAK1_cyt_ also interacts with NFR5_cyt_, but no interaction was ever observed between *Lj*BAK1_cyt_ and NFR1_cyt_. After GST pulldown, bead-bound proteins were analyzed by immunoblotting using anti-HIS and anti-GST antibodies. These experiments were repeated five times.

### *L. japonicus* PUB13 interacts with and ubiquitinates NFR5

Since *LjPUB13* is transcriptionally regulated during symbiosis and the gene is implicated in sustained nodule organogenesis, we explored the involvement of *Lj*PUB13 in Nod factor signalling. Thus, we tested the possibility of *Lj*PUB13 to interact with the Nod factor receptors NFR1 and NFR5.

First, we investigated these supposed interactions *in vitro*. The cytoplasmic regions of NFR1 or NFR5 were expressed in *E. coli* as fusions to a HIS tag and were used in an *in vitro* binding assay together with the *Lj*PUB13-GST fusion protein. We found that *Lj*PUB13 interacted strongly with NFR5_cyt_, while the *Lj*PUB13-NFR1_cyt_ interaction was usually undetectable (Fig. 5B). Sometimes a weak interaction of *Lj*PUB13-NFR1_cyt_ was observed but this result was not always reproducible, and therefore cannot be fully considered as possible.

Moreover, we tested whether *L. japonicus* BAK1, can interact with the receptors NFR1 and NFR5. *Lj*PUB13 strongly interacts with *Lj*BAK1 and it is plausible that *Lj*PUB13 acts together with *Lj*BAK1 as a complex. In addition, BAK1 is a well-known membrane co-receptor for many membrane receptor kinases (Chinchilla et al., 2009). Thus, in *in vitro* binding assays, GST-*Lj*BAK1_cyt_ was indeed found to interact with HIS-NFR5_cyt_ but no interaction was observed between GST-*Lj*BAK1_cyt_ and HIS-NFR1_cyt_ (Fig. 5B).

We verified the interaction of *Lj*PUB13 with the Nod factor receptors, using a BiFC assay in *Nicotiana benthamiana*. Since the ARM domain of PUB13 is responsible for the specificity in protein-protein interactions of E3 ligases, we created a truncated fusion protein, where only the ARM domain of *Lj*PUB13 (Fig. S2) was linked to N- or C-terminal half of YFP. We observed a strong reconstituted YFP signal on *N. benthamiana* leaf cells when *Lj*PUB13_ARM_ was co-expressed with *Lj*BAK1, NFR1 or NFR5 (Fig. 6). Two other receptor-like kinases, *Lj*CLAVATA2 (Krusell et al., 2011) and *Lj*LYS11 (Rasmussen et al., 2016), were used as negative control interactions. Indeed, no YFP signal was detected when *Lj*PUB13_ARM_ was co-expressed with either of the two control proteins. These results show that the ARM domain of *Lj*PUB13 can recognize and interacts with *L. japonicus* receptor kinases in a specific manner. In addition, the *Lj*BAK1-NFR5 interaction was also confirmed by BiFC *in planta* (Fig. 6).

**Figure 6.**
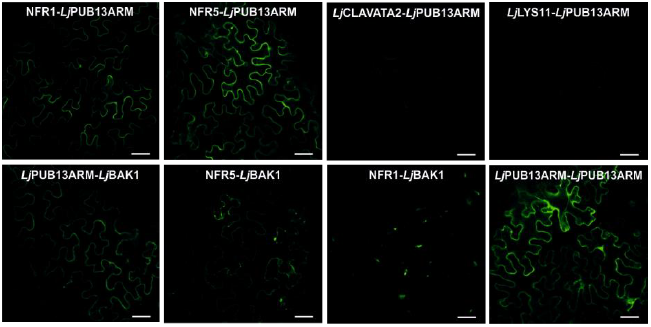
*In planta* interactions of *Lj*PUB13_ARM_ and *Lj*BAK1 with the Nod Factor receptors NFR1 and NFR5. YFP split into N- and C-terminal halves were fused to the ARM domain of *Lj*PUB13 or to full length *Lj*BAK1, NFR1 and NFR5. *N. benthamiana* leaves were cotransformed with different construct combinations and leaf epidermal cells were observed via confocal microscopy. The experiment was repeated four times with similar results. Bars, 50μm.

Finally, we examined whether *Lj*PUB13 can ubiquitinate the Nod factor receptors NFR1 and NFR5 since *Lj*PUB13 was shown to be a functional E3 ligase (Fig. 1B). The *E. coli*-produced HIS-NFR1_cyt_ or HIS-NFR5_cyt_ fusion proteins were used together with GST-*Lj*PUB13 in an *in vitro* ubiquitination assay. Interestingly, *Lj*PUB13 polyubiquitinated the cytosolic region of NFR5 and we detected high-molecular-weight proteins above the HIS-NFR5_cyt_ (Fig. 7). The NFR5 ubiquitination by *Lj*PUB13 was demonstrated by using two different E2 enzymes, UbcH6 and UbcH5b. On the contrary, when HIS-NFR1_cyt_ was tested as a substrate of *Lj*PUB13, no ubiquitination activity was observed (Fig. 7). Thus, *L. japonicus* PUB13 specifically ubiquitinates the Nod factor receptor NFR5.

**Figure 7.**
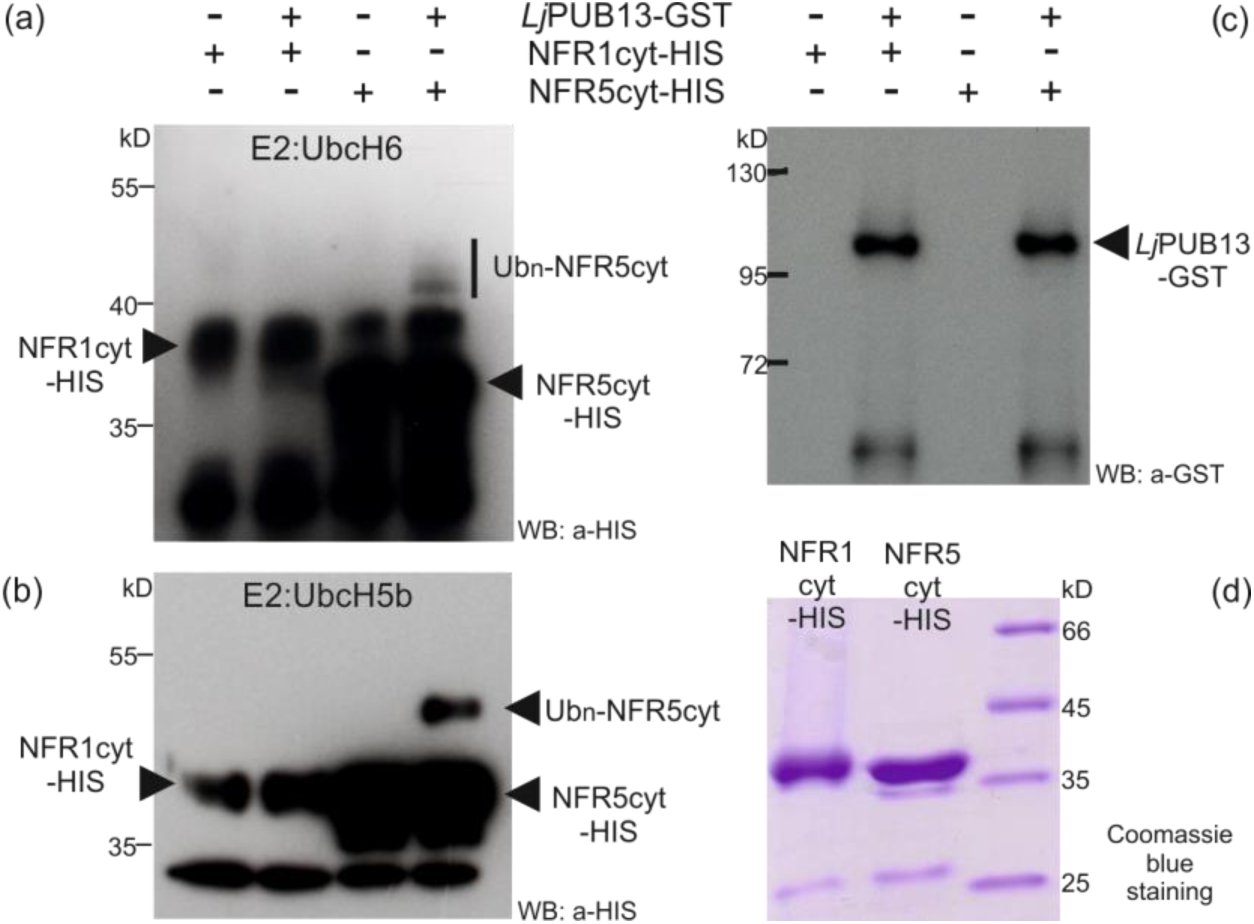
*Lj*PUB13 specifically ubiquitinates the Nod Factor receptor NFR5. *Lj*PUB13 was purified as GST fusion, while the cytoplasmic regions of NFR1 and NFR5 were purified as HIS fusion proteins. (**a, b**) The NFR5 ubiquitination was detected by an anti-HIS antibody, using two different E2 ligases: UbcH6 (a) and UbcH5b (b). No ubiquitination was observed when NFR1 was used as a substrate. **(c)** The presence of *Lj*PUB13 at the ubiquitination reactions was detected by an anti-GST antibody (input). **(d)** Purified NFR1-HIS and NFR5-HIS used at the ubiquitination reactions. The experiments were repeated three times with similar results.

## DISCUSSION

E3 ligases were identified as important signalling components in nitrogen-fixing symbiosis in model legumes (Shimomura et al., 2006; Kiss et al., 2009; Yano et al., 2009; Mbengue et al., 2010; Den Herder et al., 2012, Vernié et al., 2016). However, the targets of all the symbiotic E3 ligases remain to be identified and the signalling events downstream of their action remain unexplored. Here, we report that a PUB-ARM E3 ligase that possesses ubiquitination activity (Fig. 1) plays a direct role in Nod factor signalling in *Lotus japonicus*, through its interaction and ubiquitination of the Nod factor receptor NFR5. *Lj*PUB13 is, thus, involved in the successful establishment of *L. japonicus*-rhizobium symbiosis, likely modulating the NFR5 protein levels or activity.

We found that *L. japonicus* PUB13 is involved in the successful formation of nitrogen-fixing nodules. All three *pub13* mutant alleles displayed reduced and delayed nodule organogenetic capacity (Fig. 4). On the other hand, bacterial infection inside root hairs was normally sustained (Fig. 4D). This suggests that *LjPUB13* is required for the signalling events that lead to nodule organogenesis in the cortex, rather than the infection events. Furthermore, considering the low accumulation of *LjPUB13* gene transcripts in mature nodules (Fig. 2) and the successful colonization of nodules by rhizobium in the *pub13* mutants (Fig. S6), we envision that PUB13 E3 ligase plays a role in the initial stages of *L. japonicus*-rhizobium symbiosis establishment, rather than the later stages of nodule development and colonization.

The reduced nodulation phenotype could be attributed to the general reduced plant growth exhibited by *pub13* mutants (Fig. 3). However, the number of nodules was significantly different in *pub13* mutants from that of the wild-type plants also when expressed per unit of root length (Fig. 4B). This strongly suggests that a reduced nodulation is not directly related to a shorter root. The fact that the nodulation capacity is independent of the shoot/root length is further supported by studies in hypernodulation mutants, like *har1* (Krusell at al., 2012; Nishimura et al., 2002), where an excessive number of nodules are produced by a very short plant (Fig. S4). Moreover, RNAi knockdown of *LjnsRING* E3 ligase, a nodule-specific gene involved in the early infection and mature nodule function (Shimomura et al., 2006), and mutation of *Amsh1*, a deubiquitination enzyme (Malolepszy et al., 2015), affect plant growth in *L. japonicus*; this indicates that some proteins involved in (de) ubiquitination may act as nodes for plant growth and nodulation pathways. This is further supported by the expression of *LjPUB13* observed in developing lateral roots at non-inoculated plants (Fig. 2C). The growth defective phenotype of *L. japonicus pub13* mutants has also been reported for the *Arabidopsis pub13* mutants (Antignani et al., 2015).

The expression of *LjPUB13* observed in nodule primordia (Fig. 2), together with the findings that *Lj*PUB13 interacts with (Fig. 5, 6) and directly ubiquitinates the NFR5 receptor (Fig. 7), strongly supports the requirement for *LjPUB13* during the early stages of nodule organogenesis in *L. japonicus*-rhizobia symbiosis. The defective nodulation phenotype observed in *pub13* mutants suggests a positive regulatory role of *Lj*PUB13 in rhizobial symbiosis. On the contrary, the symbiotic E3s *Mt*PUB1 and SINA4 (Mbengue et al., 2010; Den Herder et al., 2012) exhibit a clear negative role on the early steps of the infection process. This indicates that different roles may be performed by various E3 ligases during symbiosis.

*LjPUB13* is the closest putative ortholog of *Arabidopsis PUB13* in *L. japonicus*. In *Arabidopsis*, PUB13 has been implicated in FLS2-mediated flg22 signalling (Lu et al., 2011; Zhou et al., 2015) and, recently, in LYK5-mediated chitooctaose responses (Liao et al., 2017). We show here that the defence responses downstream of flg22 perception are independent of PUB13 in the roots of this model legume. In contrast to what has been reported for *Arabidopsis* leaves (Lu et al., 2011; Zhou et al., 2015), our results from ROS production, and defence gene regulation (Fig. S7, S8) show that *LjPUB13* does not appear to be involved in flg22-dependent defence responses in *L. japonicus* roots. In line with this, our *in vitro* assays show that *Lj*PUB13 may not directly ubiquitinate *Lj*FLS2 (Fig. S9), as has been shown for its *Arabidopsis* counterpart (Lu et al., 2011). However, we cannot rule out that the involvement of *Lj*PUB13 in plant immunity may be manifested in other parts of the plant or under conditions that have not been addressed in this study.

The differences observed in plant immunity responses in *L. japonicus pub13* mutant lines compared to *Arabidopsis* mutants, could be attributed to a host-dependent specialisation of function for these PUB proteins. Functional differentiation of orthologous PUB genes was also found for *Arabidopsis PUB17* and *Brassica napus ARC1* (Yang et al., 2006). Alternatively, a redundancy in the roles of PUB proteins in *L. japonicus* could mean that paralogs may be responsible for the roles that have been assigned to PUB13 in *Arabidopsis*. Blast analyses against *L. japonicus* PUB13 in *Lotus* base v.3.0 revealed the presence of, yet uncharacterized, proteins with similarity to PUB13. Proteins with highest similarity (appr. 50% identity) are presented in Fig. S1. Future studies are needed to examine possible implication of these proteins in defence responses in *L. japonicus*.

Based on our results, we propose a plausible mechanism where *Lj*PUB13 acts on NFR5 post-translationally. Ubiquitination of NFR5 may lead to degradation, modulation of activity or re-localization. In any case, it is expected that NFR5 turnover is essential for efficient nodule organogenesis, and the recruitment of *Lj*PUB13 ensures the onset and/or continuation of NFR5-mediated signalling and the successful initiation of nodule formation.

Along this line, and although NFR5 internalization from the plasma membrane has not been shown directly as yet, an association of NFR5 with the clathrin-mediated endocytosis has been proposed (Wang et al., 2015b). A clathrin protein (CHC1) was shown to interact with the Rho-like GTPase ROP6 (Wang et al., 2015a), an interacting partner of NFR5 (Ke et al., 2012) in *L. japonicus*. Interestingly, *ROP6* silencing in roots by RNAi did not affect the rhizobium entry in root hairs, but inhibited the IT growth through the root cortex, which resulted in the development of fewer nodules per plant (Ke et al., 2012).

We also show that *Lj*PUB13 physically and specifically interacts with the cytosolic regions of *Lj*BAK1 and NFR5 *in vitro* and *in vivo*. Interestingly, the cytosolic region of *Lj*BAK1 was also shown to interact with the cytosolic region of NFR5 (Fig. 5 and 6), suggesting that PUB13/BAK1 may act as a complex in the establishment of *L. japonicus*-rhizobium symbiosis. The recruitment of *Lj*BAK1 in this case is anticipated, considering the ubiquitous role of BAK1 in interactions of multiple membrane receptors kinases (Chinchilla et al., 2009; Lu et al., 2011). In conclusion, based on the knowledge that both ubiquitination and deubiquitination of proteins play a major role in root nodule symbiosis (Shimomura et al., 2006; Kiss et al., 2009; Yano et al., 2009; Mbengue et al., 2010; Den Herder et al., 2012; Malolepszy et al., 2015; Vernié et al., 2016) and on the results presented here, we suggest that *Lj*PUB13 has a role in the establishment of the *L. japonicus*-rhizobium symbiosis and acts as a positive regulator in nodule formation through the post-transcriptional control of NFR5.

## ACKNOWLEDGEMENTS

This work was partially supported by the Postgraduate Programs 3817 & 3439 of the Department of Biochemistry and Biotechnology, University of Thessaly (to KKP). DT and SR were supported by the Danish National Research Foundation grant no. DNRF79. DT was supported by a STSM (290713-031130) from Cost Action FA1103. The authors wish to thank Prof. Jens Stougaard (Aarhus University) for scientific support and for making corrections and comments on the manuscript; and Prof. Claus Schwechheimer (Technische Universität München) for hosting DT in his laboratory by providing financial support through the SFB924 and for making comments on the manuscript. The authors thank Mickael Blaise for providing the plasmids for HIS-NFR1_cyt_ and HIS-NFR5_cyt_ protein expression, Zoltán Bozsóki for helping with the ROS assay, and Finn Pedersen for taking care of the plants in the greenhouse.

## SUPPORTING INFORMATION

**Table S1:** Primers used for cloning into expression vectors

**Table S2:** Primers used in qRT-PCR

**Table S3:** Primers used for cloning into GoldenGate and Gateway vectors

**Figure S1:** Phylogenetic tree of amino acid sequences of *Lj*PUB13 with previously characterized PUBs of other species and *L. japonicus* uncharacterized PUBs

**Figure S2:** Amino acid sequence alignment of *Lj*PUB13 with *At*PUB13

**Figure S3:** Expression levels of *Lj*PUB13 in *pub13* LORE1 mutants.

**Figure S4:** Inoculated 28-day-old wild type and homozygous *pub13.3* mutants

**Figure S5:** Formation of nodules in *pub13.1 vs. har1* mutants.

**Figure S6:** Nodule sections of wild type, *pub13.1* and *pub13.2* plants

**Figure S7:** ROS accumulation in roots of *pub13* mutants *vs* wt plants.

**Figure S8:** Expression of defence genes in *L. japonicus* wild type and *pub13* mutants

**Figure S9:** *Lj*FLS2 ubiquitination tests

